# Laboratory analysis of glucose, fructose, and sucrose contents in Japanese common beverages for the exact assessment of beverage-derived sugar intake

**DOI:** 10.1101/2021.08.13.456286

**Authors:** Yoshitaka Ando, Yoshiji Ohta, Eiji Munetsuna, Hiroya Yamada, Yuki Nouchi, Itsuki Kageyama, Genki Mizuno, Mirai Yamazaki, Ryosuke Fujii, Hiroaki Ishikawa, Koji Suzuki, Koji Ohashi

**Affiliations:** Department of Informative Clinical Medicine, Fujita Health University School of Medical Sciences, Toyoake, Aichi, Japan; Department of Chemistry, Fujita Health University School of Medicine, Toyoake, Aichi, Japan; Department of Biochemistry, Fujita Health University School of Medicine, Toyoake, Aichi, Japan; Department of Hygiene, Fujita Health University School of Medicine, Toyoake, Aichi, Japan; Department of Joint Research Laboratory of Clinical Medicine, Fujita Health University Hospital, Toyoake, Aichi, Japan; Department of Medical Technology, Kagawa Prefectural University of Health Sciences, Takamatsu, Kagawa, Japan; Department of Preventive Medical Sciences, Fujita Health University School of Medical Sciences, Toyoake, Aichi, Japan

**Author notes:** Corresponding author: (KO), (EM).

**Keywords:** Glucose, Fructose, Sucrose, sugar content, Japanese common beverages, Adverse health effects

## Abstract

**Background:** The adverse health effects of sugar-sweetened beverage consumption have been studied worldwide. There are several reports on actual sugar contents in sugar-sweetened beverages. However, there is no recent report on actual sugar contents in Japanese sugar-sweetened beverages. Therefore, we attempted to analyze glucose, fructose, and sucrose contents in Japanese common beverages.

**Methods:** Glucose, fructose, and sucrose contents in 49 beverages including 8 energy drinks, 11 sodas, 4 fruit juices, 7 probiotic drinks, 4 sports drinks, 5 coffee drinks, 6 green tea drinks, and 4 tea drinks were determined using the enzymatic methods.

**Results:** Tow zero calorie drinks, 2 sugarless coffee drinks, and 6 green tea drinks contained no sugar. Three coffee drinks contained only sucrose. The orders of median glucose, fructose, and sucrose contents in categorized beverages containing sugars were as follows: for glucose, fruit juice > energy drink ≥ soda >> probiotic drink > black tea drink > sports drink; for fructose, probiotic drink ≥ energy drink > fruit juice > soda >> sports drink > black tea drink; and for sucrose, black tea drink > energy drink ≥ probiotic drink > fruit juice > soda > coffee drink >> sports drink. The rate of total fructose content in total sugar content in 38 sugar-containing beverages was approximately 40-60%. The total sugar content analyzed was not always equivalent to carbohydrate content indicated on the nutrition label.

**Conclusions:** These results indicate that actual sugar content in Japanese common beverages is necessary for the exact assessment of beverage-derived sugar intake.

## Introduction

Sugar-sweetened beverages (SSBs) are consumed together with fruit juice and milk worldwide [1]. Therefore, the adverse effect of SSB consumption on health has been examined all over the world. A cross-national analysis of 75 countries on the relationship of soft drink consumption to global overweight, obesity and, diabetes has shown that soft drink consumption is significantly linked to overweight, obesity, and diabetes prevalence worldwide [2]. Furthermore, overconsumption of SSBs has been reported to be associated with an increased risk of various metabolic diseases such as metabolic syndrome [3, 4], type 2 diabetes [5, 6], nonalcoholic fatty liver disease (NAFLD) [7], cardiovascular disease [8], hypertension [9, 10], and hyperuricemia [11, 12], and with the risk of overall cancer [13]. Especially, the adverse effect of overconsumed SSBs containing fructose and/or high-fructose corn syrup (HFCS) has been reported to be associated with the risk of obesity [14, 15], NAFLD [16–18], type 2 diabetes [14], hyperuricemia [15, 19], cardiometabolic disease [14, 20], and hypertension [15, 20]. Recently, it has been suggested that fructose intake during pregnancy has an adverse impact on the next generation [21–27]. However, actual fructose intake from SSBs containing fructose and/or HFCS has not been estimated based on the actual content of fructose in beverages including SSBs which are commonly consumed.

Until now, analysis of the actual contents of sugars such as glucose, fructose, and sucrose in SSBs made with and without HFCS which are commonly consumed has been performed in USA and Poland. In 2011, Ventura et al. [28] reported the content and composition of glucose, fructose, and sucrose in 23 popular sweetened beverages made with HFCS alone, which were purchased from retailers in East Los Angeles, CA, USA, with focus on fructose content. The authors found that the total sugar content in popular SSBs varied from the information provided by the manufacture/vendor with some having more and some having less sugar than the label and that the type of sugar listed on the label of beverages was not always consistent with the type of sugar detected in the corresponding beverages. Furthermore, the authors suggested that the tendency for use of HFCS containing higher fructose could be contributing to higher fructose consumption than would be assumed. In 2014, Walker et al. [29] reported the contents of glucose, fructose, galactose, lactose, maltose, and sucrose in 14 sodas as the beverage and 19 fruit juices, which were purchased from retailers in East Los Angeles, CA, USA, and analyzed using three different methods in three independent laboratories. The authors showed that the actual amount of free fructose in some sodas was higher than the expected amount, that popular sodas contained 50% more fructose than glucose, although all sodas contained little galactose, lactose, and maltose, while a few sodas contained sucrose, and that some fruit juices had twice as much fructose as glucose. In 2014, Bilek et al. [30] reported the analysis of glucose, fructose, and sucrose contents in 15 mineral and spring water-based beverages, which were purchased from shops at the area of Rzeszow, Poland. The authors showed that mineral and spring water-based beverages contained more glucose and fructose than sucrose, although the contents of glucose and fructose were almost equal and that the total content of glucose, fructose, and sucrose in each mineral and spring water-based beverage was almost consistent with the content of total carbohydrate in the corresponding beverage which was declared by the manufactures. The authors suggested that the excessive consumption of sugars such as glucose, fructose, and sucrose from mineral and spring water-based beverages might be disadvantage for human health. In 2015, White et al. [31] reported the contents and composition of glucose, fructose, and higher saccharides such as maltose, maltotriose, and malototetraose, which were analyzed in 2 independent laboratories, in 80 carbonated beverages sweetened with HFS-55, which were randomly collected from retail stores in USA. The authors showed that fructose and glucose comprised 55.58% and 39.71% of total sugars in 80 carbonated beverages sweetened with HFS-55, respectively and that the total content of sugars in the carbonated beverages was in close agreement with published specifications in industry technical data sheets, published literature values, and governmental standards and requirements. In 2017, Varsamis et al. [32] analyzed total glucose content and total fructose content in 4 soft drinks consumed commonly in Australia, Europe, and USA, and showed that both total glucose content and total fructose content in 4 soft drinks were different among Australia, Europe, and USA. Besides, a cross-sectional survey of the amounts of free sugars and calories in carbonated SSBs on sale in the UK has demonstrated that the content of free sugars in carbonated SSBs on sale in the UK, which is estimated from the product packaging and nutrient information panels of the beverages, is high and that a high free sugar content in carbonated SSBs is a major contributor to free sugar intake [33].

There is a report on the actual contents of several sugars in Japanese common beverages including SSBs. In 2011, Inukai et al. [34] reported the analysis of glucose and sucrose contents in 45 Japanese popular soft drinks. i.e., 9 carbonated drinks, 3 carbonated diet drinks, 3 energy drinks, 11 sports drinks, 3 tea drinks, 4 juice drinks, 4 100% juices, 5 coffee drinks, and 3 lactic acid drinks (i.e., probiotic drinks), using a biochemistry analyzer. The authors showed that glucose and sucrose contents in all soft drinks were widely variable, although glucose content was high in energy drinks, carbonated soft drinks, lactic acid drinks, juice drinks, and 100% juices, while sucrose content was high in lactic acid drinks, tea drinks, coffee drinks, energy drinks, and 100% juices. In addition, it has been demonstrated that SSB consumption assessed using a food frequency questionnaire is associated with an increased risk of type 2 diabetes in Japanese adults without prior history of diabetes [35], a middle-aged Japanese population [36], and Japanese adults with impaired glucose tolerance [37]. However, the association of SSB consumption with an increased risk of type 2 diabetes has not been evaluated based on the assessment of the actual contents of sugars in Japanese common beverages including SSBs yet. Besides, there is no available information on the actual contents of sugars such as glucose, fructose, and sucrose in Japanese common beverages on sale at present.

Therefore, we attempted to analyze actual glucose, fructose, and sucrose contents in 49 Japanese common beverages on sale at present, which included 8 energy drinks, 11 sodas, 4 fruit juices, 7 probiotic drinks, 4 sports drinks, 5 coffee drinks, 6 green tea drinks, and 4 black tea drinks, using the enzymatic methods for the exact assessment of beverage-derived sugar intake in a Japanese population.

## Materials and methods

### Chemicals

Glucose, fructose, and sucrose used as the standard were purchased from Fujifilm Wako Pure Chemical Co. (Osaka, Japan). These sugars were the highest reagent grade. Other chemicals were also obtained from Fujifilm Wako Pure Chemical Co. (Osaka, Japan). All chemicals were used without further purification. Glucose oxidase (GOD) and invertase were purchased from Sigma-Aldrich Japan (Tokyo, Japan). Phosphoglucose isomerase was purchased from NIPRO (Osaka, Japan).

### Samples of Japanese beverages used for sugar analysis

The following 48 Japanese beverages were purchased from several supermarkets in Toyoake and Nagoya, Aichi, Japan: 8 energy drinks, 11 sodas, 4 fruit juices, 7 probiotic drinks, 4 sports drinks, 5 coffee drinks, 6 green tea drinks, and 4 black tea drinks. The information on carbohydrate content in each beverage was obtained from the declared nutrition label of each beverage.

### Measurement of sugars in beverages

For the measurement of glucose and fructose in beverages, each beverage was diluted with 40 volumes of distilled water. For the measurement of sucrose content in beverages, each beverage was diluted with 40 volumes of 50 mM phthalate buffer (pH 4.0) containing invertase (0.02 U/μL).

Glucose, fructose, and sucrose in beverages were measured by the enzymatic methods reported by Al-Mhanna et al. [38] with a slight modification. Glucose content in beverages was measured using a commercial kit, EKDIA XL ‘EIKEN’ GLUII (EIKEN CHEMICAL Co., Ltd., Tokyo, Japan). In brief, a mixture of 25 μL of Reagent 1 (ATP2Na) and 4 μL of 40-fold diluted beverage sample was transferred into a 96 well multiplate and incubated at 37°C for 30 min. Then, 100 μL of Reagent 2 including hexokinase (HK), glucose 6-phospahte dehydrogenase, and NADP^+^ was added to the incubated medium. The mixture medium was incubated at 37°C for 30 min. After incubation, the absorbance of the reaction medium was measured at 340 nm for NADPH formed by reduction of NADP^+^, using a multilabel plate reader, 2330 ARVO X (PerkinElmer Inc., Waltham, MA, USA). Fructose measurement in beverages was conducted as follows: fructose in beverage samples was converted to fructose 6-phosphate in the presence of ATP by HK. The formed fructose 6-phosphate was converted to glucose 6-phosphate by PGI. Before HK treatment, glucose present in beverages was removed by conversion of glucose to gluconolactone by GOD. The formed glucose 6-phosphate was converted to 6-phosphogluconate in the presence of NADP^+^ by glucose 6-phospahte dehydrogenase at which time NADP^+^ was reduced to NADPH. Accordingly, the above-described enzymatic method for glucose measurement in beverages was used for fructose measurement in beverages using Reagent 1 including GOD (1 U/μL) and Reagent 2 including PGI (0.02 U/μL) under the same reaction condition as used for the glucose measurement. The absorbance of the reaction medium was measured at 340 nm using the same plate reader as used for the glucose measurement. Fructose concentration in beverages was calculated using the concentration of glucose converted from fructose. For sucrose measurement in beverages, the above-described diluted beverage samples containing invertase were incubated at 20°C for 90 min at which time sucrose was degraded to glucose and fructose. Total glucose (after hydrolysis) in beverages was measured using the above-described glucose method. The measurement of fructose in the degraded sucrose was not performed to analyze sucrose content in beverages easily and rapidly because glucose content is equal to fructose content in sucrose. Sucrose content in beverages was calculated using the following formula: [Sucrose] = 2{[glucose] Total (after hydrolysis) – [glucose] Initial (before hydrolysis)}. The contents of glucose, fructose, and sucrose in beverages were calculated using standard curves made with authentic D-glucose, D-fructose, and sucrose, respectively.

Glucose, fructose, and sucrose contents in each beverage are expressed as the mean value of at least duplicate or triplicate determinations. Glucose, fructose, and sucrose contents and their total content in each categorized beverage are expressed as the median value with the range of each sugar content. Total fructose content in beverages is expressed as the sum amount of fructose itself, i.e., free fructose, and sucrose-derived fructose.

## Results

### Glucose, fructose, and sucrose contents in beverages

The contents of glucose, fructose, and sucrose in 49 beverages that are commonly consumed at present in Japan were analyzed using the enzymatic methods. All beverages analyzed were classified into 8 beverage categories, i.e., energy drink, soda, fruit juice. probiotic drink, sports drink, coffee drink, green tea drink, and black tea drink. As shown in Table 1, 1 energy drink (Zero Calorie), 2 sodas (Zero Calorie), 2 coffee drinks (Sugarless), and all 6 green tea drinks did not contain any glucose, fructose, and sucrose, while 7 energy drinks, 9 sodas, 4 fruit juices, 7 probiotic drinks, 4 sports drinks, 3 coffee drinks, and 4 black tea drinks contained glucose, fructose and/or sucrose. Of 39 sugar-containing beverages analyzed, Oronamin, Coca Cola Energy, and Monster Energy in the category of energy drink had the highest glucose content (74.1 g/L), the highest fructose content (58.9 g/L), and the highest sucrose content (81.0 g/L), respectively. Most of beverages in the categories of energy drink, soda, fruit juice, probiotic drink, and black tea drink contained glucose, fructose, and/or sucrose in a considerably high amount, although glucose, fructose and/or sucrose contents in those beverages were variable.

**Table 1.**
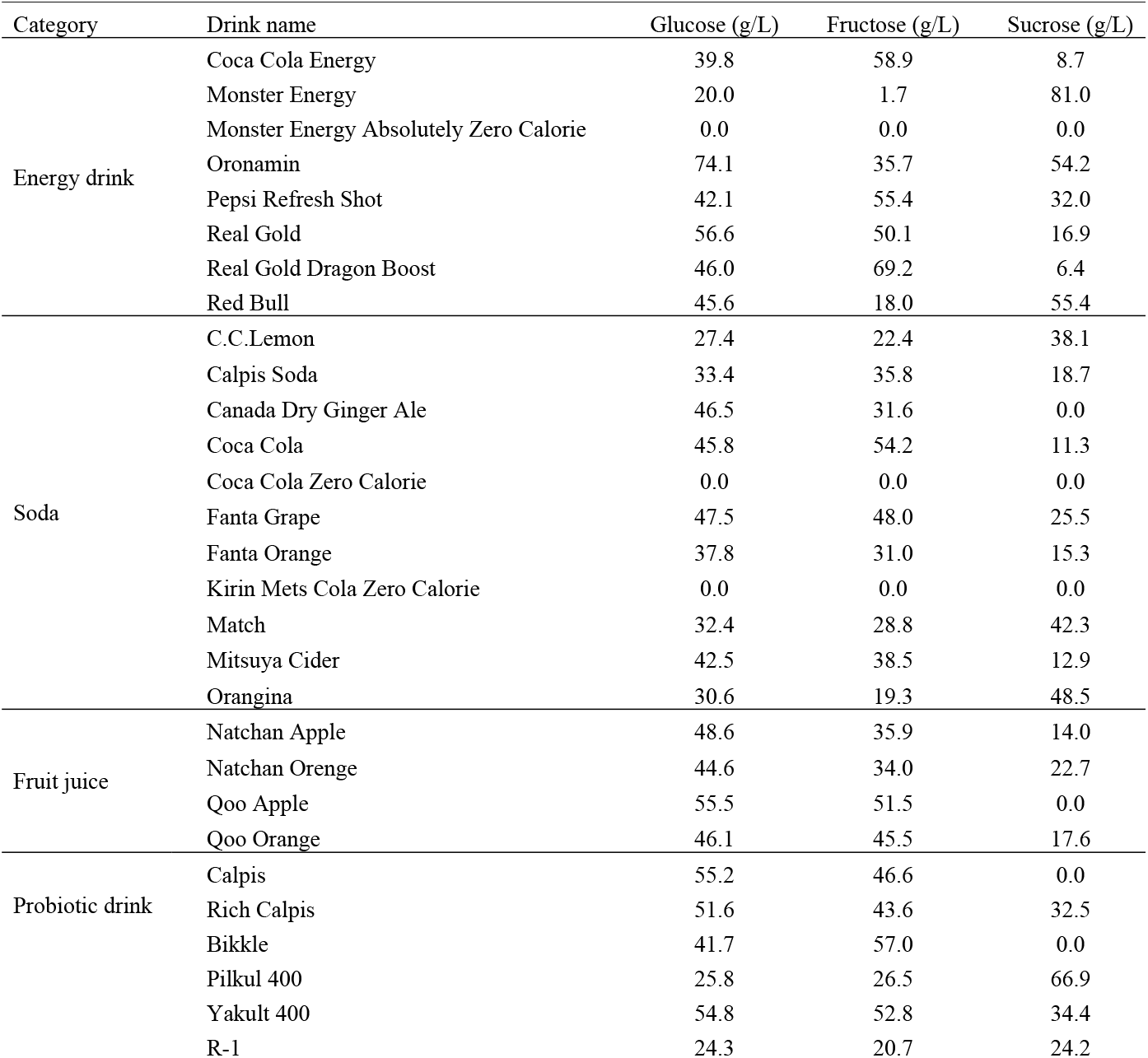
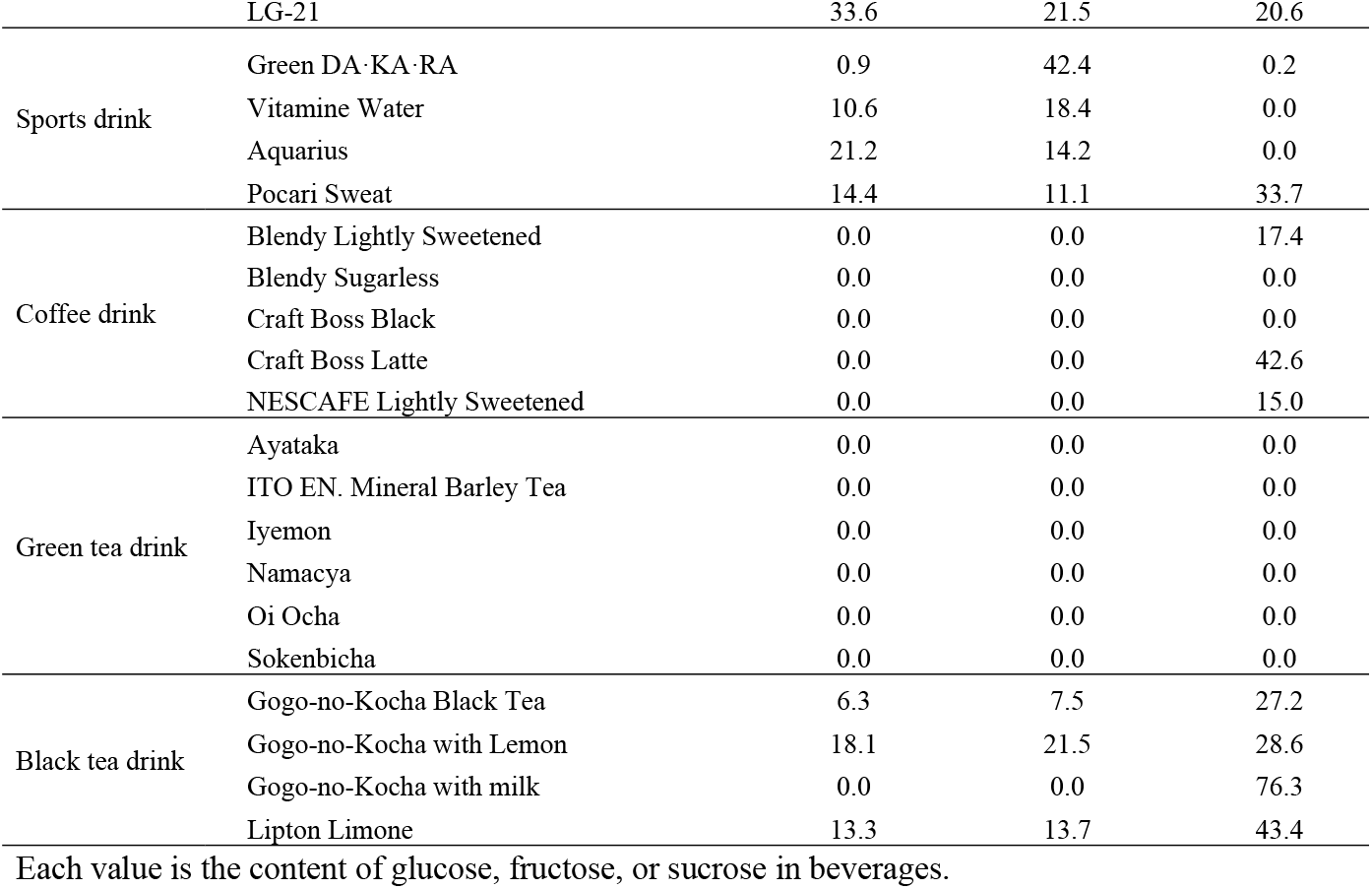
Glucose, fructose, and sucrose contents in 48 Japanese popular beverages.

### Glucose, fructose, and sucrose contents and total sugar contents in categorized beverages

Table 2 shows the median contents of glucose, fructose, sucrose, and their total and the ranges of each sugar and total sugar content in 7 categorized beverages containing sugars. When glucose, fructose, and sucrose contents in categorized beverages containing sugars were compared based on their median contents, the order of each median sugar content was as follows: for median glucose content in 6 categorized beverages, fruit juice > energy drink ≥ soda >> probiotic drink > black tea drink > sports drink; for median fructose content in 6 categorized beverages, probiotic drink ≥ energy drink > fruit juice > soda >> sports drink > black tea drink; and for median sucrose content in 7 categorized beverages, black tea drink > energy drink ≥ probiotic drink >> fruit juice > soda > coffee drink >> sports drink. The order of median total sugar content in 7 categorized beverages containing sugars was as follows: energy drink > fruit juice ≥ probiotic drink > soda > black tea drink >> sports drink > coffee drink. In 7 categorized beverages, sports drink and coffee drink had lower median sugar content than energy drink, fruit juice, probiotic drink, soda, and black tea drink. In addition, the range of each sugar content and total sugar content in categorized beverages containing sugars was widely variable (Table 2).

**Table 2.** Median contents of glucose, fructose, and sucrose and their total and ranges of each sugar and total sugar contents in 7 categories of Japanese common beverages.

| Category | Categorized number | Glucose (g/L) | Fructose (g/L) | Sucrose (g/L) | Total Sugar (g/L) |
| --- | --- | --- | --- | --- | --- |
| Energy drink | 7 | 43.8<br>(20.0-74.1) | 42.9<br>(1.7-69.2) | 24.5<br>(6.4-81.0) | 120.3<br>(102.8-164.0) |
| Soda | 9 | 40.8<br>(27.4-47.5) | 31.0<br>(19.3-54.2) | 15.3<br>(11.3-48.5) | 87.9<br>(78.0-121.0) |
| Fruit juice | 4 | 47.3<br>(44.6-55.5) | 40.7<br>(34.0-51.5) | 15.8<br>(17.6-22.7) | 104.1<br>(98.5-109.1) |
| Probiotic drink | 7 | 27.0<br>(24.3-55.2) | 43.6<br>(20.7-57.0) | 24.2<br>(20.6-66.9) | 101.7<br>(69.2-142.0) |
| Sports drink | 4 | 13.1<br>(0.9-21.2) | 16.3<br>(11.1-42.4) | 0.2<br>(0.0-33.7) | 39.5<br>(29.0-59.2) |
| Coffee drink | 5 | 0.0<br>(0.0) | 0.0<br>(0.0) | 15.0<br>(0.0-42.6) | 15.0<br>(0.0-42.6) |
| Black tea drink | 4 | 15.6<br>(0.0-18.1) | 10.6<br>(0.0-21.5) | 36.0<br>(27.2-76.3) | 69.3<br>(41.1-76.3) |

The ranges of each sugar and total sugar contents in categorized beverages are shown in parenthesis.

### Rates of glucose, fructose, and sucrose contents to total sugar content in beverages

As shown Fig. 1A, the rate of glucose content in the total content of glucose, fructose, and sucrose in 33 beverages except 4 beverages containing little or no glucose was approximately15-60% and the mean rate was 36.1%. As shown Fig. 1B, the rate of fructose content to the total content of glucose, fructose, and sucrose in 33 beverages except 4 beverages containing little or no fructose was approximately16-98% and the mean rate was 38.1%. As shown Fig. 1C, the rate of sucrose content to the total content of glucose, fructose, and sucrose in 31 beverages except 7 beverages containing little or no sucrose was approximately16-98% and the mean rate was 41.1%. Thus, the mean rate of each sugar content in sugar-containing beverages was almost similar, although the rate of each sugar content itself was widely variable in individual beverages.

**Fig. 1.**
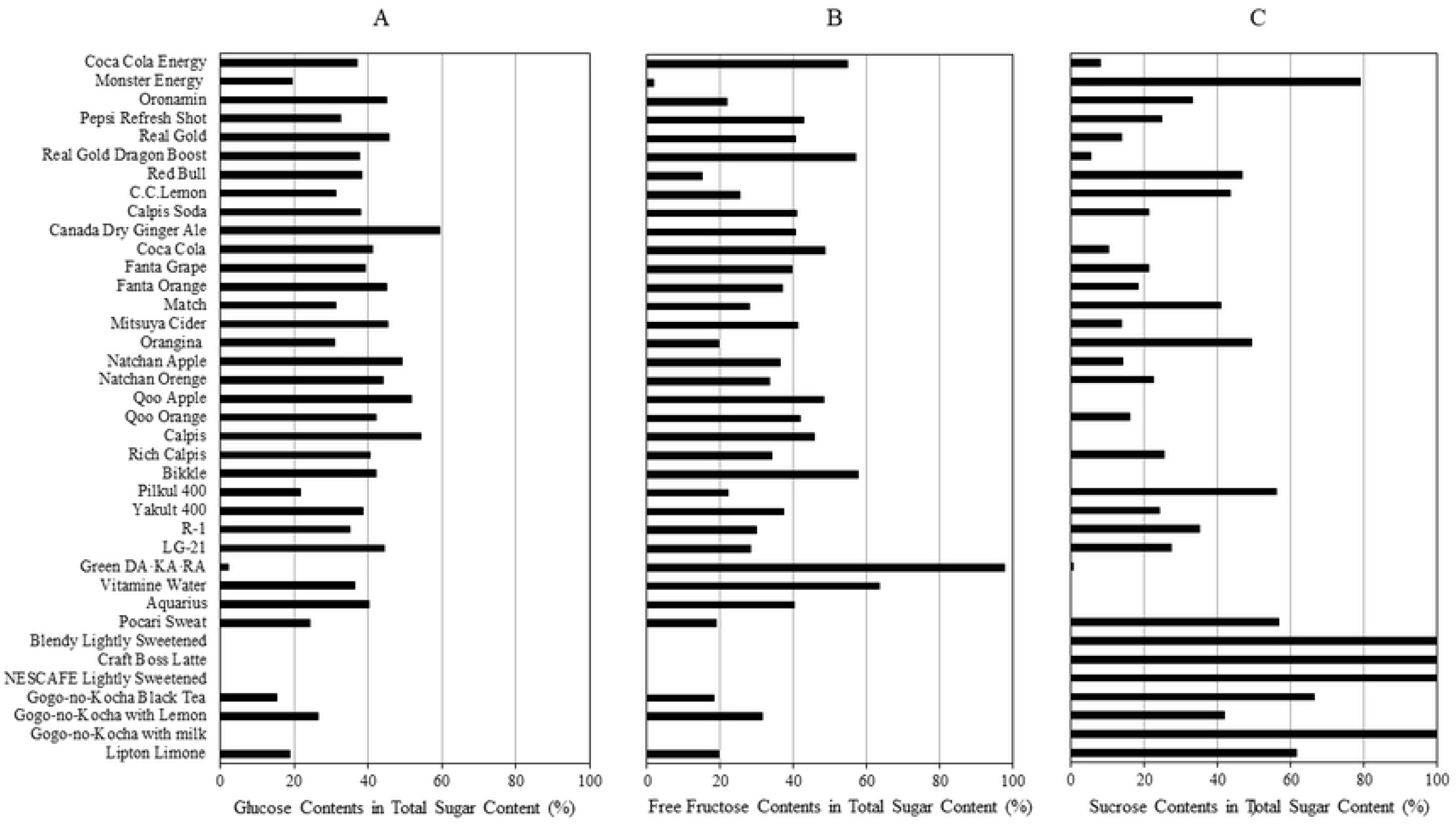
Rates of glucose, fructose, and sucrose contents to total sugar content in 38 Japanese common beverages. All beverages containing glucose, fructose and/or sucrose listed in Table 1 were analyzed. The rates of glucose, fructose, and sucrose contents in each beverage were calculated from the ratio of each sugar content to the total content of glucose, fructose, and sucrose in the corresponding beverage.

### Rate of total fructose content to total sugar content in beverages

Figure 2 shows the rate of the total content of fructose and fructose-derived from sucrose in the total contents of glucose, fructose, and sucrose in 38 beverages except 13 beverages not containing fructose and/or sucrose which were included in the categories of energy drink, soda, coffee drink, and green tea drink. The rate of the total fructose content in 37 beverages was approximately 40-60% of the total sugar content, although that rate of one drink, Green DA□KA□RA in the category of sports drink, was almost 100%. (Fig. 2).

**Fig. 2.**
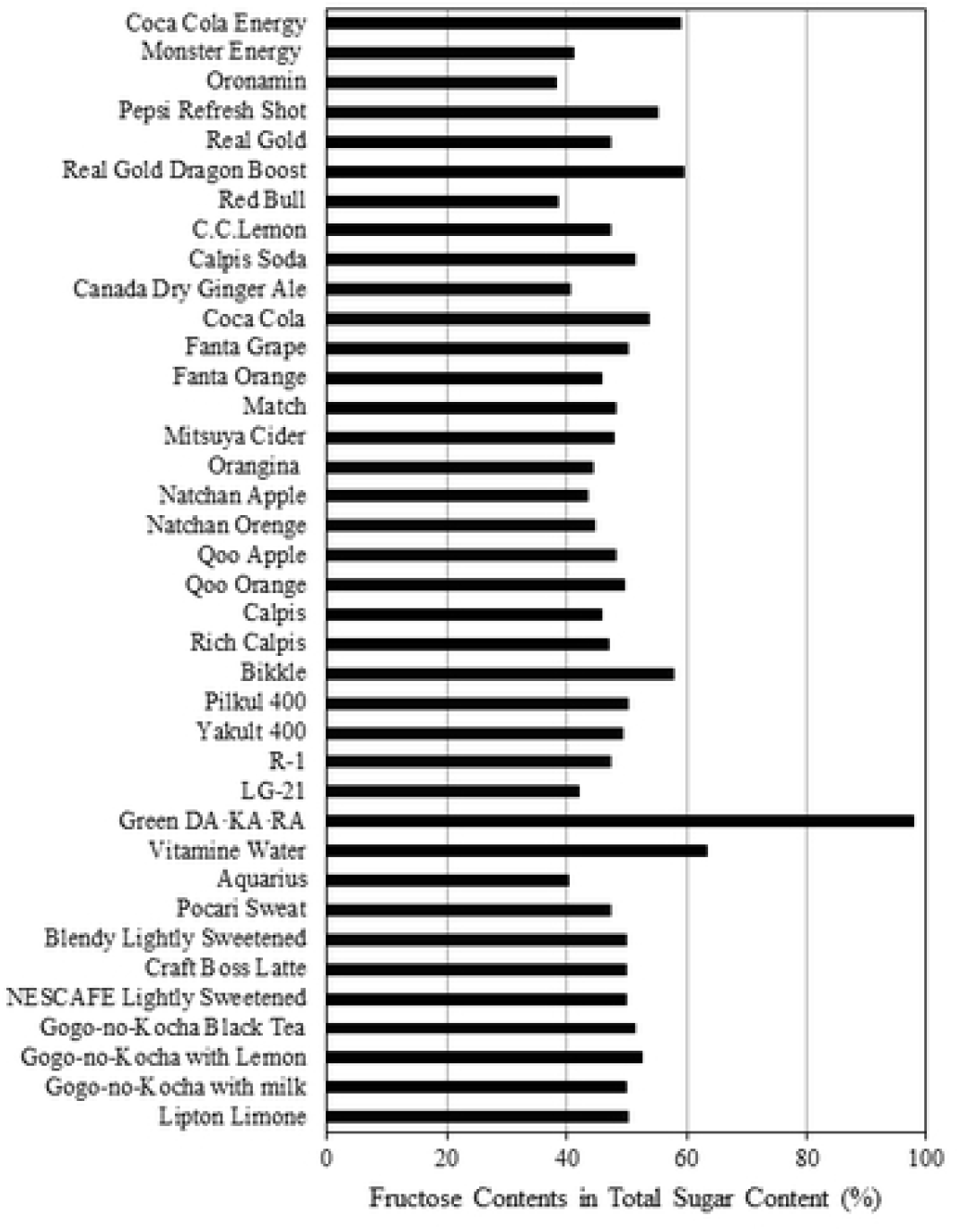
Rate of total fructose content to total sugar content in 36 Japanese common beverages. All beverages containing fructose and/or sucrose listed in Table 1 were analyzed. Total fructose content is expressed as the sum of fructose content and a half of sucrose content because sucrose consists of one molecular glucose and one molecular fructose. The rate of total fructose content in each beverage was calculated from the percentage of total fructose content in the total content of glucose, fructose, and sucrose in the corresponding beverage.

### Comparison between the total content of sugars analyzed and carbohydrate content indicated on the nutritional label in each beverage

The total content of glucose, fructose, and sucrose analyzed in 49 beverages was compared to carbohydrate content indicated on the nutrition label in those beverages. As shown in Table 3, the total content of three sugars analyzed was considerably well consistent with carbohydrate content indicated on the nutrition label in 43 beverages. However, there was a large difference between the total sugar content analyzed and carbohydrate content indicated on the nutrition label in five drinks, i.e., Monster Energy, Fanta Grape, Pilkul 400, Yakult 400, LG-2, and NESCAFE Lightly Sweetened (Table 3). The difference between the total sugar content and carbohydrate content in the three drinks was above 20 g/L (Table 3).

**Table 3.**
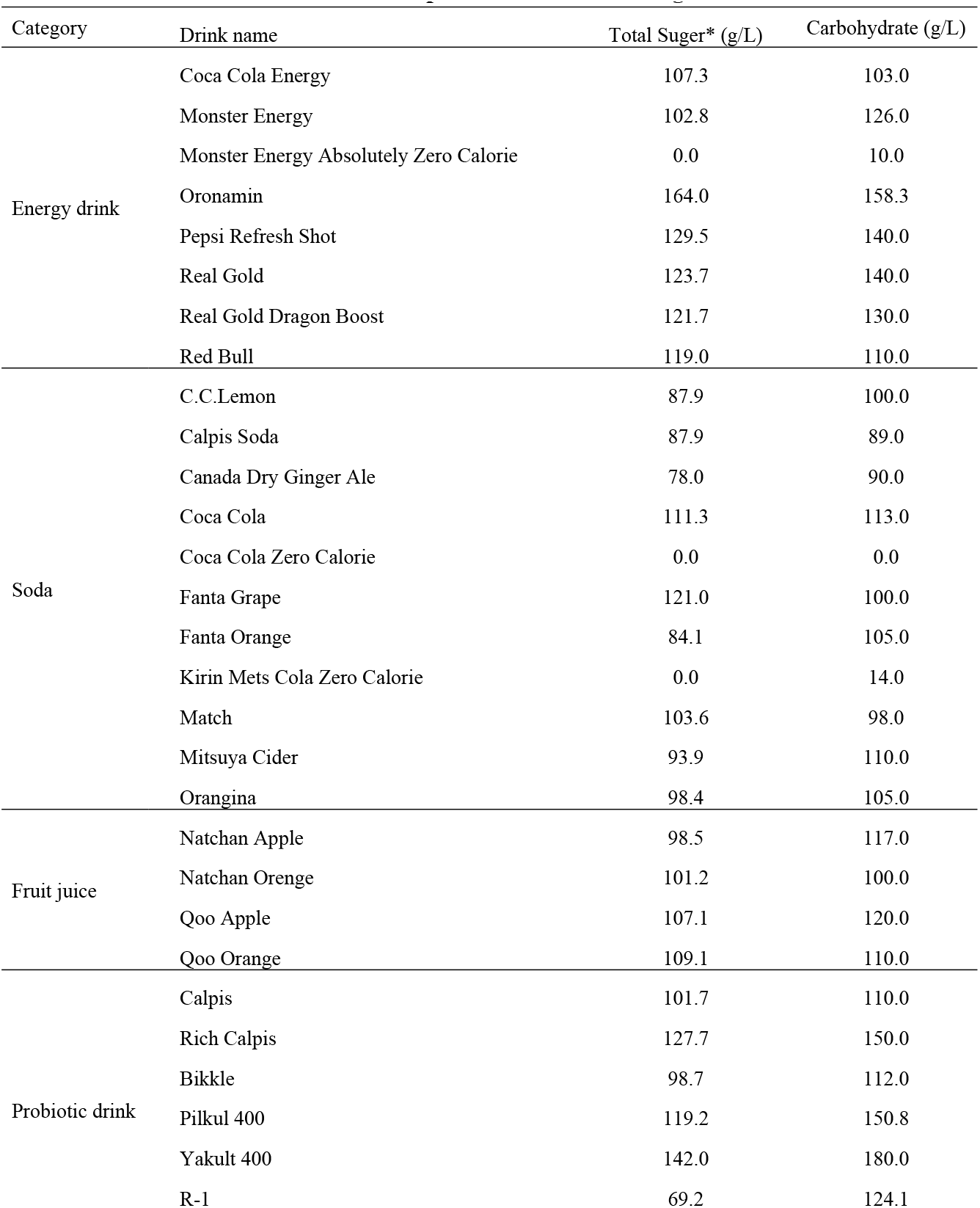
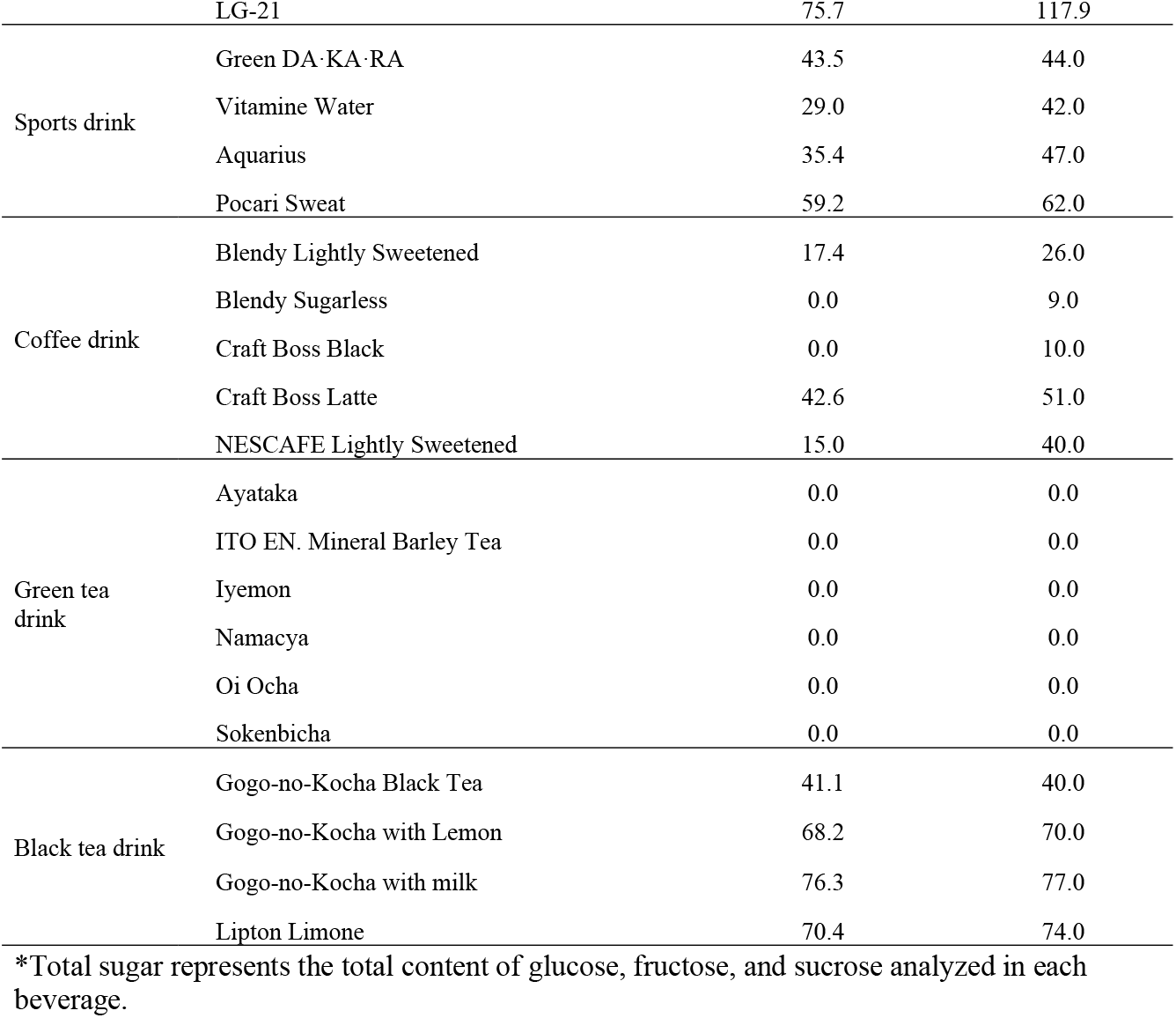
Comparison between total sugar content analyzed and carbohydrate content indicated on the nutrition label in 49 Japanese common beverages.

## Discussion

Beverages including SBBs have been consumed with fruit juices and milk commonly worldwide [1]. Analysis of actual sugar content in common beverages including SBBs has been performed in several countries including USA, Poland, and Japan. The analysis performed in these countries has shown that most of SBBs contain several sugars such as glucose, fructose, and sucrose [28–31, 34, 39]. However, there is a report showing that when total glucose content and total fructose content are analyzed in 4 soft drinks, which are consumed commonly in Australia, Europe, and USA are different among Australia, Europe, and USA [32]. Bilek et al. [30] have suggested based on the analysis of glucose, fructose, and sucrose contents in 15 mineral and spring water-based beverages on sale in Poland that the excessive consumption of sugars such as glucose, fructose, and sucrose from mineral and spring water-based beverages might be disadvantage for human health. In addition, it has been reported worldwide that fructose overconsumption from beverages causes the risk of obesity [14, 15], NAFLD [16–18], type 2 diabetes [14], hyperuricemia [15, 19], cardiometabolic disease [14, 20], and hypertension [15, 20]. In Japan, there are some reports showing that SSB consumption assessed using a food frequency questionnaire is associated with an increased risk of type 2 diabetes in a Japanese population [35–37]. However, the association of SSB consumption with an increased risk of type 2 diabetes in a Japanese population has not been evaluated based on the assessment of the actual content of sugars in Japanese common beverages including SSBs. In addition, there is no available information on the actual content of sugars such as glucose, fructose, and sucrose in Japanese common beverages on sale at present.

In the present study, we analyzed the actual content of glucose, fructose, and sucrose in 48 beverages including 8 energy drinks, 11 sodas, 4 fruit juices, 7 probiotic drinks, 4 sports drinks, 5 coffee drinks, 6 green tea drinks, and 4 black tea drinks that are commonly consumed at present in Japan, using the enzymatic methods, for the exact assessment of beverage-derived sugar intake in a Japanese population. Of 49 beverages analyzed, 1 energy drink (Zero Calorie), 2 sodas (Zero Calorie), 2 coffee drinks (Sugarless), and 6 green tea drinks did not contain any glucose, fructose, and sucrose. Thus, calorie free drinks or sugarless drinks in Japanese common beverages on sale at present were found to contain no sugar. Of the remaining 38 beverages, i.e., 7 energy drinks, 9 sodas, 4 fruit juices, 7 probiotic drinks, 4 sports drinks, 3 coffee drinks, and 4 black tea drinks were found to contain glucose, fructose and/or sucrose. Most of Japanese common beverages in the categories of energy drink, soda, fruit juice, probiotic drink, and black tea drink were found to contain considerably high amounts of glucose, fructose, and/or sucrose. As to the actual contents of glucose, fructose, and/or sucrose in Japanese common beverages, similar results have been reported previously [34]. In addition, we analyzed glucose, fructose, and sucrose contents in one oolong tea drink, Suntory Oolong tea. As a result, the oolong drink did not contain any sugar as found in green tea drinks, although this data is not shown in Table 1. Thus, no glucose, fructose, and sucrose were detected in 3 calorie-free drinks and 9 sugarless drinks in the categories of energy drink, soda, coffee drink, green tea drink, and oolong tea drink. Accordingly, calorie-free drinks and sugarless drinks seem to be recommended to reduce the overconsumption of sugar-containing beverages such as SSBs on sale in Japan.

The median glucose, fructose, and sucrose contents and the median total sugar content in 7 categorized beverages containing sugars, i.e., energy drink, soda, fruit juice, probiotic drink, sports drink, coffee drink, and black tea drink, were further analyzed. When the median glucose and fructose contents were compared among 6 categorized beverages except categorized coffee drink not containing glucose and fructose, the median glucose content was higher in fruit juice, energy drink, and soda than in probiotic drink, black tea drink, and sports drink. The median fructose content was higher in probiotic drink, energy drink, fruit juice, and soda than in sports drink and black tea drink. The median sucrose content in 7 categorized beverages was much lower in sport drink than in other categorized beverages. In addition, the range of each sugar content in 7 categorized beverages containing sugars was widely variable. Thus, the contents of glucose, fructose, and sucrose in 7 categorized beverages containing sugars were found to be largely depending on the composition of three sugars. The median total sugar content in 7 categorized beverages was higher in energy drink, fruit juice, probiotic drink, and soda than in black tea drink, sports drink, and coffee drink. The range of total sugar content in 7 categorized beverages was widely variable. These results suggest that the exact assessment of sugar intake from Japanese common beverages should be performed based on the actual content of sugars in each beverage.

Of sugar components in sugar-containing beverages such as SSB, fructose is associated with health problem closely because overconsumption of fructose from beverages causes an increased risk of obesity [14, 15], NAFLD [16–18], type 2 diabetes [14], hyperuricemia [15, 19], cardiometabolic disease [14, 20], and hypertension [15, 20]. In the present study, the total content of fructose itself and sucrose-derived fructose in 38 Japanese common beverages containing sugars was found to be approximately 40-60% of the total content of glucose, fructose, and sucrose in those beverages, although the total fructose content in one drink in the category of sports drink was almost 100%. Thus, most of Japanese common beverages containing sugars were found to contain fructose and sucrose-derived fructose in a considerable high rate. This result suggests that fructose overconsumption from Japanese common beverages should be estimated based on the actual content of fructose and sucrose-derived fructose in those beverages.

Ventura et al. [28] have shown in 23 popular SSBs on sale in USA that the total content of sugars i.e., glucose, fructose, and sucrose varies from the information provided by the manufacture/vendor and that the type of sugar listed on the label is not always consistent with the type of sugar detected. In contrast, Bilek et al. [30]have shown in 15 mineral and spring water-based beverages on sale in Poland that the total content of glucose, fructose, and sucrose is almost consistent with total carbohydrate content which is declared by the manufactures. White et al. [31] have also shown in 80 carbonated beverages sweetened with HFS-55 on sale in USA that the total content of sugars, i.e., glucose, fructose, and higher saccharides is in close agreement with published specifications in industry technical data sheets, published literature values, and governmental standards and requirements. Although the analysis of sugar content in Japanese common beverages has been reported previously [34], however, such a comparison between the total content of sugars analyzed and carbohydrate content indicated on the nutrition label or the food label provided by the manufacture/vendor has not been performed in Japanese common beverages until now. In the present study, the total content of glucose, fructose, and sucrose, which were analyzed by the enzymatic method, was found to be considerably well consistent with carbohydrate content indicated on the nutrition label in 43 Japanese common beverages used as the sample. However, the total content of glucose, fructose, and sucrose analyzed was largely different from carbohydrate content indicated on the nutrition label in 6 beverages The difference between the total sugar content and carbohydrate content in these beverages was above 20 g/L. However, the cause of the difference between both contents in these drinks is unclear at present This result indicates that there is a case in which the exact amounts of sugars consumed from Japanese common beverages cannot be assessed from carbohydrate content indicated on the nutrition label correctly. Therefore, it can be thought that the actual content of sugars in beverages is useful for the exact assessment of sugar intake from beverages in the case of consumed Japanese common beverages.

## Conclusion

The results obtained in the present study indicate that the actual content of glucose, fructose, and sucrose analyzed and the rate of each sugar content to total sugar content in Japanese common beverages vary widely depending on the composition of sugars in each beverage, that the rate of total fructose content in Japanese common beverages containing sugars is approximately 40-60%, and that the total content of sugars analyzed is considerably well consistent with carbohydrate content indicated on the nutrition label in most of Japanese common beverages but there are beverages showing a large difference between both contents. These results also indicate that the actual content of sugars in Japanese common beverages is necessary for the exact assessment of beverage-derived sugar intake in a Japanese population.

## Acknowledgments

This work was supported by the Ministry of Education, Culture, Sports, Science, and Technology of Japan via a Grant-in-Aid for Scientific Research and Grant-in-Aid, 2020-2023 (No. 20H04134).

## Author Contributions

Conceptualization: Yoshitaka Ando, Yoshiji Ohta, Eiji Munetsuna, Hiroya Yamada, Genki Mizuno, Mirai Yamazaki, Hiroaki Ishikawa, Koji Ohashi.

Formal analysis: Yoshitaka Ando, Yoshiji Ohta, Eiji Munetsuna, Hiroya Yamada, Mirai Yamazaki, Ryosuke Fujii, Koji Suzuki.

Funding acquisition: Hiroya Yamada.

Investigation: Yoshitaka Ando, Yuki Nouchi, Itsuki Kageyama, Genki Mizuno.

Methodology: Yoshitaka Ando, Hiroaki Ishikawa, Koji Ohashi.

Writing – original draft: Yoshitaka Ando, Yoshiji Ohta, Eiji Munetsuna, Hiroya Yamada, Koji Ohashi.

Writing – review & editing: Yoshitaka Ando, Yoshiji Ohta, Koji Ohashi.

